# Transcriptomic data support a nocturnal bottleneck in the ancestor to gecko lizards

**DOI:** 10.1101/592873

**Authors:** Brendan J. Pinto, Stuart V. Nielsen, Tony Gamble

## Introduction

Behavioral shifts between different light environments, such as changes from diurnal to nocturnal activity patterns, have led to major modifications to the vertebrate eye over evolutionary time (Walls 1934; Walls 1942; Davies et al. 2012). These modifications involve changes to the types of photoreceptor cell present in the retina, changes in cell morphology, and alterations to the ancestral gene complement used by these cells to transmit light into a biochemical signal (Walls 1942; Simões et al. 2015; Lamb and Hunt 2017). Two types of photoreceptors are present in most vertebrate retinas, rods and cones, used for low-light vision and daylight vision, respectively (Kojima et al. 1992; Lamb 2013). Rods and cones possess significant differences in their sensitivities to light (rods being more light-sensitive than cones) and phototransduction speed (cones transmit biochemical signals faster than rods) (Li et al. 2010), providing tradeoffs in the selective forces driving adaptation to differing light environments (Simões et al. 2015; Schott et al. 2016). Indeed, lineages that have experienced dramatic evolutionary shifts in light environment during their evolution, such as snakes and geckos, have seen concomitant changes photoreceptor cell complement, resulting in photoreceptor cell loss and subsequent ‘transmutation’, a process where cones take on a rod-like morphology or rods take on a cone-like morphology (Walls 1934; Walls 1942; Pedler and Tilley 1964; Tansley 1964; Underwood 1970; Zhang et al. 2006; Schott et al. 2016).

The ancestral tetrapod eye utilized five photopigments: four opsins in cone cells: LWS, RH2, SWS1, and SWS2; and one opsin in rod cells: RH1 (Okano et al. 1992; Davies et al. 2012; Lamb and Hunt 2017). However, many vertebrate groups lack this ancestral visual opsin complement due to subsequent adaptations to nocturnal, fossorial, and other low-light aquatic environments, including: crocodilians (Emerling 2017a), burrowing rodents (Emerling and Springer 2014), snakes (Davies et al. 2009; Simões et al. 2015), whales (Levenson and Dizon 2003), and geckos (Crescitelli et al. 1977; Kojima et al. 1992; Yokoyama and Blow 2001). Characterizing the presence and absence of components of phototransduction signaling pathway – particularly photopigments and key members of the phototransduction cascade – among extant species, in a phylogenetic context, facilitates investigation of visual adaptation to a particular light environment (Serb and Oakley 2005). For example, investigating the loss of SWS2 and RH2 photopigments in most mammals provided evidence of a so-called “nocturnal bottleneck”, a period of dim-light adaptation early in their evolutionary history (Walls 1942; Menaker et al. 1997; Gerkema et al. 2013).

Geckos are thought to be ancestrally nocturnal – with multiple, independent transitions back to diurnality throughout their evolutionary history – making them an important model for investigating how changes in light environment impact vision (Walls 1934; Walls 1942; Kojima et al. 1992; Röll 2000; Roth & Kelber 2004; Gamble et al. 2015). All gecko species appear to have a retina composed of a single photoreceptor type having, at least superficially, a rod-like morphology (Walls 1942; Underwood 1951; Underwood 1954; Röll 2000). In fact, it was the presence of rod-like cells in the retinas of the limbless pygopodids that provided important evidence that these lizards were, in fact, geckos (Underwood 1954; Underwood 1957). Examination of tokay gecko (*Gekko gecko*) opsins and other phototransduction genes have shown that, despite their rod-like morphology, they produce solely cone proteins, consistent with the ‘transmutation’ hypothesis (Walls 1942; Crescitelli et al. 1977; Kojima et al. 1992; Yokoyama and Blow 2001; Zhang et al. 2006). Furthermore, detailed examination of the cellular ultrastructure reveals many characteristics unique to cones and the cellular morphology is only superficially rod-like (Röll 2000). Despite their historic importance for studying visual system evolution, nearly all studies of the molecular components of the gecko visual system were performed within the genus *Gekko* (mostly *Gekko gecko*) and a few species of Malagasy day geckos (*Phelsuma* ssp.) (Crescitelli et al. 1977; Kojima et al. 1992; Loew et al. 1994; Taniguchi et al. 1999; Taniguchi et al. 2001; Yokoyama and Blow 2001; Roth et al. 2009; Liu et al. 2015). Both genera are in the family Gekkonidae, which is nested within the infraorder Gekkota (composed of seven families) and fail to show whether these changes are gecko-wide or specific only to the family Gekkonidae (Fig. 1). Indeed, Underwood (1954) suggested that transmutation occurred up to three times in geckos and gecko rod-like retinas evolved repeatedly through convergent evolution. More recently, the examination of pseudogenes in the *Gekko japonicus* genome suggested a step-wise loss of visual opsins, with loss of SWS2 occurring approximately ~202 million years ago (mya), preceding the loss of the rod opsin, RH1, ~81 mya (Emerling 2017b), well after the hypothesized divergence of extant gekkotan families ~120 mya (Gamble et al. 2011; Gamble et al. 2015). It remains unclear whether the cone-to-rod transmutation and loss of rod photoreceptors occurred prior to the diversification of extant geckos, and, thus, is ubiquitous across all gecko species.

**Figure 1:**
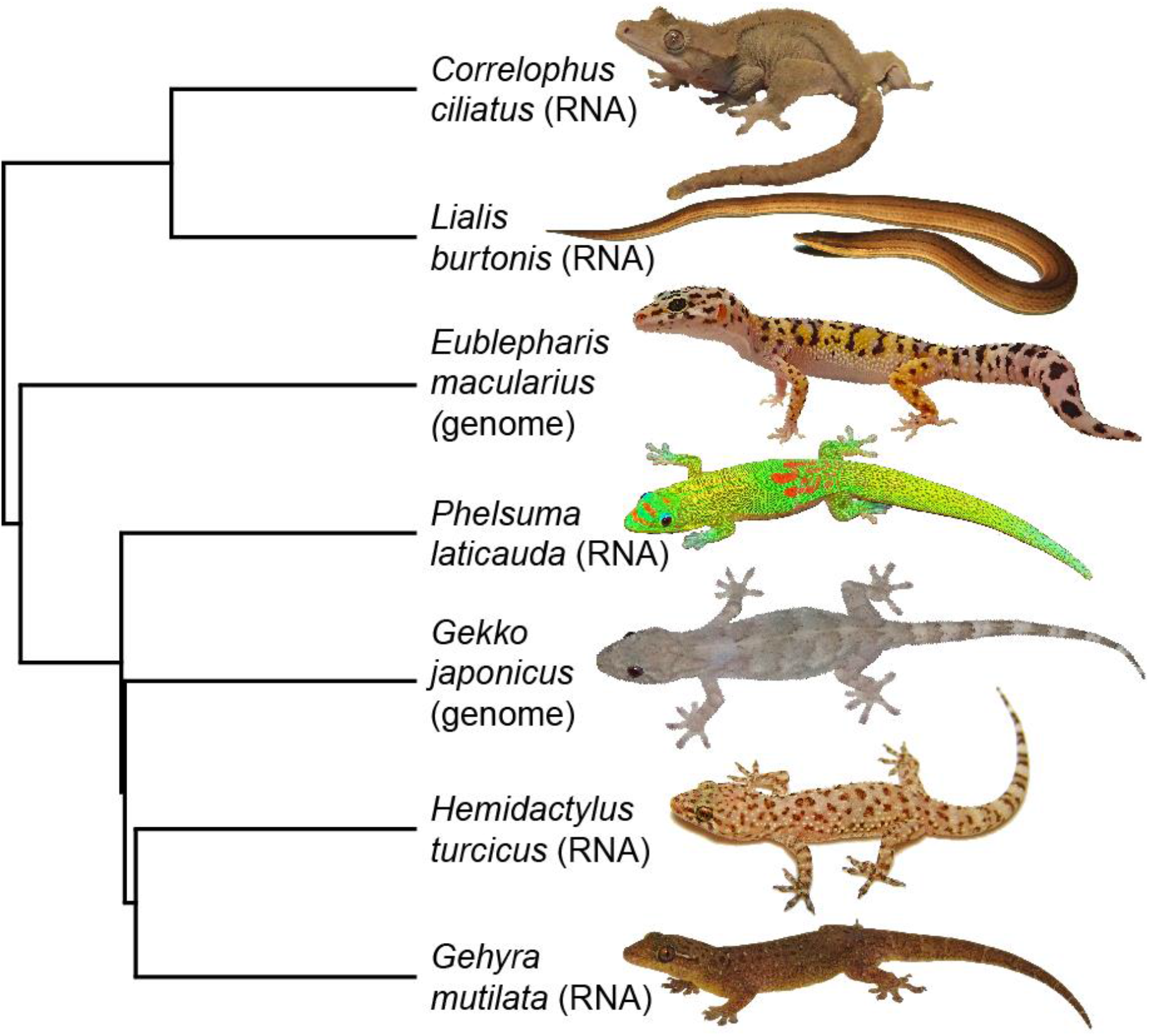
(A) Phylogeny, pruned from Gamble et al. (2015), depicting relationships among sampled gecko species (*Correlophus ciliatus*, family: Diplodactylidae; *Lialis burtonis*, family Pygopodidae; *Eublepharis macularius*, family Eublepharidae; and *Phelsuma laticauda, Gekko japonicus, Hemidactylus turcicus,* and *Gehyra mutilata*, family Gekkonidae). Data from transcriptomes or genomes are indicated.

By combining data from two published gecko genomes with six de novo eye transcriptomes, we here, for the first time, investigate the early evolution of gecko vision across the entire phylogenetic breadth of extant geckos. We find that a suite of rod genes, as well as the cone-opsin, SWS1, are not expressed in any sampled gecko species, consistent with (i) a single loss of rods and (ii) a cone-to-rod transmutation during a “nocturnal bottleneck” in the most recent common ancestor (MRCA) of extant geckos.

## Materials and Methods

### RNAseq and Transcriptome Assembly

We euthanized and removed whole eyes from 5 geckos, representing the breadth of extant gecko diversity (Fig. 1; *Correlophus ciliatus*, *Gehyra mutilata*, *Hemidactylus turcicus*, *Lialis burtonis*, and *Phelsuma laticauda*). All species are nocturnal or crepuscular except *P. laticauda*, which is diurnal. Tissues were flash frozen at −80°C in TRIzol™ reagent. RNA extraction, library prep, and transcriptome assembly processes are identical to those described by Pinto et al. (2019). Briefly, we extracted RNA using the Qiagen RNeasy™ Mini Kit and prepared RNAseq libraries with KAPA^®^ Stranded mRNA-Seq Kit (KR0960 [v5.17]). Libraries were sequenced on an Illumina^®^ HiSeq 2500 (paired-end 125 bp reads). We assembled *de novo* transcriptomes for each species using the *De novo* RNA-Seq Assembly Pipeline (DRAP) [v1.91] (Cabau et al. 2017), which is a compilation of automated assembly (Trinity [v2.4.0]; Grabherr et al. 2011) and quality-control tools to reduce transcript redundancy.

### Ortholog Identification and Phylogenetic Analyses

We downloaded a set of 35 key phototransduction genes, including opsins, assumed to be present in the ancestor to all tetrapods (Schott et al. 2018), for nine species (Supplemental Table 3) from Ensembl [v91.0]. We used BLAST, implemented in Geneious^®^ [v11.1.2] (Altschul et al. 1990; Kearse et al. 2012) to identify orthologs to these genes from annotated CDS’s from the published genomes of nine additional species, including two additional geckos (*Eublepharis macularius* and *Gekko japonicus*), and transcriptomes from a chameleon (*Chamaeleo calyptratus*; Pinto et al. 2019) and the five gecko species described above (Supplemental Table 3).

Four transcripts, GNAT2 in *Correlophus*, GNGT2 and GUCY2D in *Lialis*, and SAG in *Hemidactylus*, were not found in the assembled transcriptomes. However, since the numbers of assembled *de novo* transcripts can vary greatly when assembled from short-reads (Zhao et al. 2011), we suspected that these ‘missing’ transcripts were sequenced, just not assembled. We confirmed their presence by mapping quality-filtered RNAseq reads to the respective transcript of their closest sampled relative using Geneious^®^ [v11.1.2]. Similarly, two genes (GNAT1 and GUCY2F) had functional copies present in the two gecko genomes but were not assembled in any of the five gecko transcriptomes. We mapped RNAseq reads to GNAT1 and GUCY2F CDSs from the *G. japonicus* genome but recovered no transcript for either gene in any sampled geckos. We performed the same read mapping strategy to all rod-specific transcripts that were consistently missing from all gecko transcriptomes. To visualize these data, we produced a character matrix indicating presence/absence of each phototransduction gene from the genome or transcriptome for every sampled species (Fig. 2) using *phytools* [v0.6-60] (Revell 2012) in R (R Core Team 2008).

**Figure 2:**
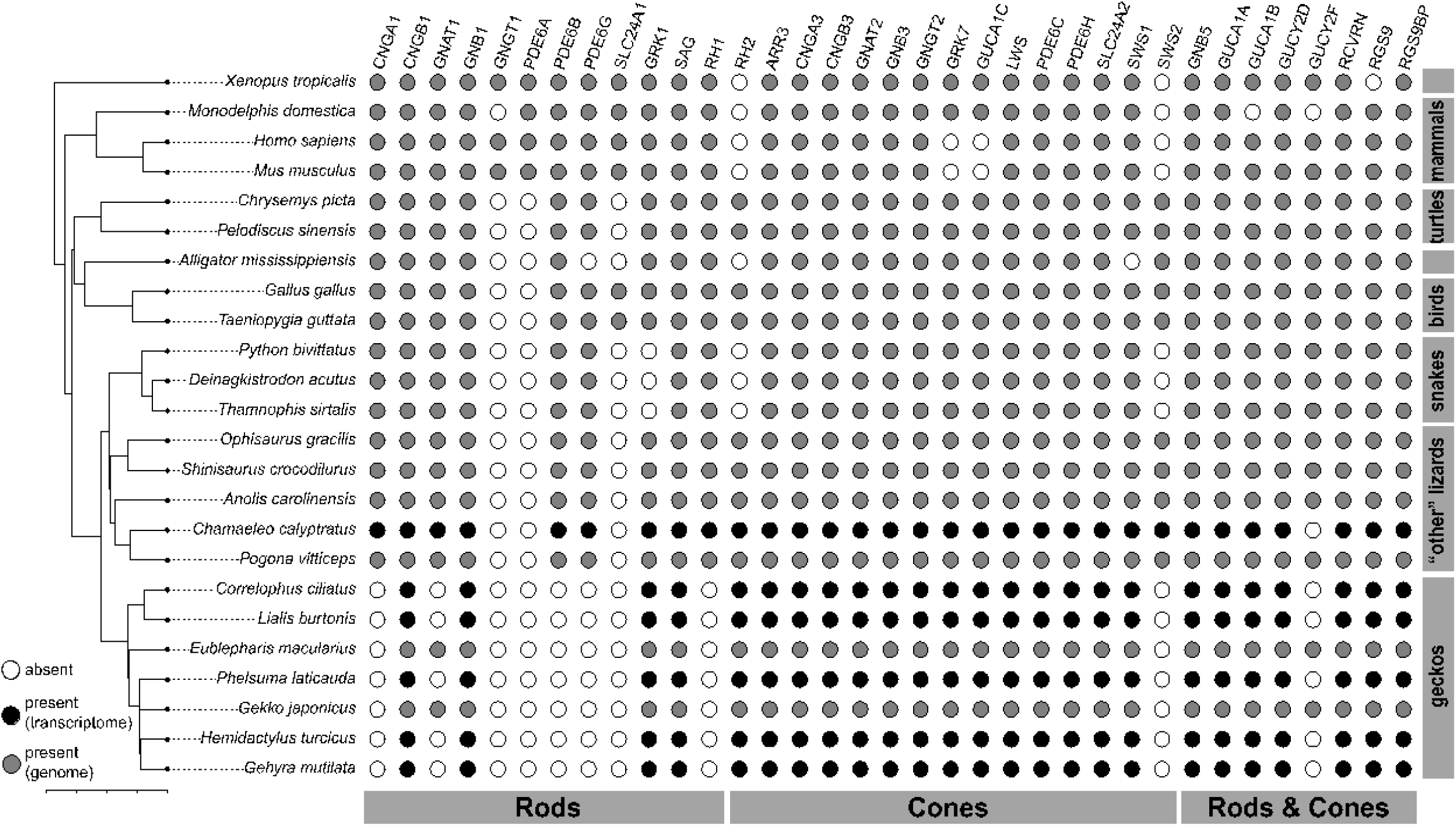
Presence/absence of 35 ancestral phototransduction genes for sampled tetrapod species. Filled gray circles indicate presence in the sampled genome, filled black circles indicate presence in the sampled transcriptome, white circles indicate absence. Expression in rods, cones, or both cell types is indicated along the bottom of the matrix. Phylogeny is a composite from Gamble et al. (2015) – geckos – and Irisarri et.al. (2017) – other vertebrates.

The difficulty assigning orthology from BLAST alone prompted us to use phylogenies to ascertain orthology for several gene families. Sequences were translation aligned using MAFFT [v7.388] implemented in Geneious^®^ [v11.1.2] (Katoh et al. 2002; Kearse et al. 2012). We generated gene trees for a subset of gene alignments (Supplemental Fig. 1–4) on the CIPRES portal (Miller et al. 2010) using the RAxML Blackbox [v8.2.10] (Stamatakis, 2014). CNGA3 sequences were constrained as monophyletic – following Lamb & Hunt (2017).

## Results and Discussion

We assembled *de novo* eye transcriptomes for five gecko species and one chameleon, *Chamaeleo calyptratus* (assembly statistics and benchmarking information are in Supplemental Table 1). We recovered the same 25 (out of 35) phototransduction genes in the RNAseq data from the five gecko eyes. We recovered 31 (out of 35) phototransduction genes from the chameleon transcriptome (Fig. 2). Eight rod-specific genes, including the visual opsin RH1, were missing in all the gecko transcriptomes, which supports the hypothesis that rod cells were lost in the MRCA of extant geckos (Fig. 1b). Similarly, the cone-specific opsin, SWS2, was missing from all sampled geckos but present in chameleon. Maximum-likelihood phylogenies from visual opsins (Supplemental Fig. 1) and several other phototransduction genes were largely concordant with previously published gene trees (Lamb and Hunt 2017) and allowed us to confirm orthology of sequences initially identified via BLAST.

While there was broad concordance between our transcriptomic data and the genome data from the geckos *Eublepharis* and *Gekko*, there were two genes that were not expressed in the eye but still had functional copies in the genomes. Transcriptomic data indicates loss of expression of the rod-specific GNAT1, which retained functional copies in the *Gekko* and *Eublepharis* genomes, suggesting an additional function for this gene outside of the eye. Similarly, the rod and cone gene, GUCY2F, was not expressed in gecko or chameleon eyes although functional copies were found in all sampled squamate genomes. Given that expression is also missing in snakes (Schott et al. 2018), the loss of GUCY2F expression in the eye is likely squamate-wide.

We observed no differences in the occurrence of core phototransduction transcripts among the sampled gecko eyes (Fig. 2). These results are consistent with the MRCA of geckos going through a nocturnal bottleneck that resulted in the transmutation of cones into a rod-like morphology and loss of rod cells (Fig. 1b). Nocturnal bottlenecks are known from mammals and crocodilians and both led to significant changes to the expression of photopigments and phototransduction genes in the eye (Gerkema et al. 2013; Emerling et al. 2017a). Snakes are also thought to have gone through a visual bottleneck, likely due to a fossorial ancestor, which, coupled with more recent transitions in diel activity, has led to repeated transmutations of rod-like and cone-like cells (Walls 1942; Schott et al. 2016; Simões et al. 2016). Thus, within squamate reptiles, two visual bottlenecks (fossoriality in snakes and nocturnality in geckos) accompanied a different set of phototransduction gene losses and subsequent cellular transmutations. In some snakes, all-cone retinas have been observed via a rod-to-cone transmutation (Schott et al. 2016). However, in geckos, the loss of RH1 and accompanying rod phototransduction genes provides conclusive evidence of a single loss of ancestral rod cells and a transmutation of ancestral cones to a rod-like morphology (Walls 1942).

Results presented here provide the first molecular evidence that gecko species spanning the gekkotan phylogeny share a visual system shaped by adaptation to a low light environment and nocturnality. Gecko eyes are characterized by the loss of the cone-opsin SWS1, rod photoreceptors, and the majority of rod-specific phototransduction genes. Prior characterizations of the gekkonid visual system using *Gekko gecko* and *Phelsuma* are thus representative of gekkotan eyes overall, at least at the molecular level (Crescitelli et al. 1977; Kojima et al. 1992; Loew et al. 1994; Taniguchi et al. 1999; Taniguchi et al. 2001; Yokoyama and Blow 2001; Zhang et al. 2006; Roth et al. 2009; Liu et al. 2015). While most rod-specific phototransduction genes that we searched for were no longer expressed in the eye, some functional rod-specific genes remained. These may be genes that retain some function elsewhere in the retina, or may have been co-opted in the transmutation of cone cells into their rod-like morphology. Further research that localizes expression of these remaining rod-specific genes in the gekkotan retina would help resolve this.

Additional lines of evidence also support a nocturnal ancestor in geckos. These traits are widespread in geckos and include numerous adaptations to a low-light lifestyle, including: sustained locomotion at low temperatures (Autumn et al. 1999); olfactory specialization (Schwenk 1993); widespread acoustic communication (Gans and Maderson 1973; Marcellini 1977); and additional eye modifications such as increased size, pupils capable of extreme constriction and dilation, and retinas lacking foveae (Röll 2001). Finally, comparative phylogenetic analyses of diel activity patterns of extant geckos also indicate the MRCA of extant geckos was nocturnal (Gamble et al. 2015). When combined with the molecular data presented here, these multiple lines of evidence overwhelmingly support a “nocturnal bottleneck” of the MRCA of extant gecko lizards. This led not only to a dramatic restructuring of the eye to adapt to low-light vision, but also to changes in nearly all aspects of gekkotan morphology, physiology, and behavior. Thus, if geckos weren’t already interesting enough, the re-evolution of diurnality has occurred repeatedly in this clade (Gamble et al. 2015). With luck, future work will elucidate the molecular evolution of visual opsins and other phototransduction genes as they adapt from ancestral nocturnality to diurnality across the breadth of independently-evolved shifts in diel activity pattern.

## Acknowledgements and funding information

We thank: M. Sea, C. Del Angel, S. Keating, R. Laver, and D. Zarkower for field assistance; C. Siler and A. Fenwick for additional samples. Hawaii Permit numbers: EX-18-02 & EX-18-06. MU IACUC AR-298; AR279; AR288. Funding from MU startup funds to T.G. and NSF-DEB1657662.

## Data Availability

SRA project for data generated in this study is available in Supplemental Table 1. Assembled transcriptomes and alignments are available via Figshare (doi:TBD).

## Author Contributions

BJP and TG developed project aims, assessed gene orthology, and wrote the manuscript. BJP assembled and annotated transcriptomes. SVN extracted RNA and made sequencing libraries. All authors conducted fieldwork.

## References

Altschul SF, Gish W, Miller W, Myers EW, Lipman DJ. 1990. Basic local alignment search tool. J Mol Biol. 215(3):403–410.

Autumn K, Jindrich D, DeNardo D, Mueller R. 1999. Locomotor performance at low temperature and the evolution of nocturnality in geckos. Evolution. 53:580–599.

Cabau C, Escudié F, Djari A, Guiguen Y, Bobe J, Klopp C. 2017. Compacting and correcting Trinity and Oases RNA-Seq *de novo* assemblies. PeerJ. 5:e2988.

Crescitelli F, Dartnall HJ, Loev ER. 1977. The gecko visual pigments: a microspectrophotometric study. J Physiol. 268:559–573.

Davies WL, Cowing JA, Bowmaker JK, Carvalho LS, Gower DJ, Hunt DM. 2009. Shedding light on serpent sight: the visual pigments of henophidian snakes. J Neurosci. 29(23):7519–7525.

Davies WL, Collin SP, Hunt DM. 2012. Molecular ecology and adaptation of visual photopigments in craniates. Mol Ecol. 21(13):3121–3158.

Emerling CA. 2017a. Archelosaurian color vision, parietal eye loss and the crocodylian nocturnal bottleneck. Mol Biol Evol. 34(3):666–676.

Emerling CA. 2017b. Genomic regression of claw keratin, taste receptor and light-associated genes provides insights into biology and evolutionary origins of snakes. Mol Phylogenet Evol. 115:40–49.

Emerling CA, Springer MS. 2014. Eyes underground: regression of visual protein networks in subterranean mammals. Mol Phylogenet Evol. 78:260–270.

Gamble T, Bauer AM, Colli GR, Greenbaum E, Jackman TR, Vitt LJ, Simons AM. 2011. Coming to America: Multiple origins of new world geckos. J Evol Biol. 24:231–244.

Gamble T, Greenbaum E, Jackman TR, Bauer AM. 2015. Into the light: Diurnality has evolved multiple times in geckos. Biol J Linnean Soc. 115:896–910.

Gans C, Maderson PFA. 1973. Sound producing mechanisms in recent reptiles: Review and comment. Amer Zool. 13:1195–1203.

Gerkema MP, Davies WI, Foster RG, Menaker M, Hut RA. 2013. The nocturnal bottleneck and the evolution of activity patterns in mammals. Proc R Soc Lond. 280(1765):20130508.

Grabherr MG, Haas BJ, Yassour M, Levin JZ, Thompson DA, Amit I, Adiconis X, Fan L, Raychowdhury R, Zeng Q, Chen Z, et al. 2011. Full-length transcriptome assembly from RNA-Seq data without a reference genome. Nature Biotechnol. 29(7):644–652.

Irisarri I, Baurain D, Brinkmann H, Delsuc F, Sire J, Kupfer A, Peterson J, et al. (2017). Phylotranscriptomic consolidation of the jawed vertebrate timetree. Nat Ecol Evol. 1(9):1370–1378.

Katoh K, Misawa K, Kuma K, Miyata T. 2002. MAFFT: a novel method for rapid multiple sequence alignment based on fast Fourier transform. Nucleic Acids Res. 30(14):3059–3066.

Kearse M, Moir R, Wilson A, Stones-Havas S, Cheung M, Sturrock S, Buxton S, Cooper A, Markoqitz S, Duran, C, et al. (2012). Geneious Basic: An integrated and extendable desktop software platform for the organization and analysis of sequence data. Bioinformatics. 28(12):1647–1649.

Kojima D, Okano T, Fukada Y, Shichida Y, Yoshizawa T, Ebrey TG. 1992. Cone visual pigments are present in gecko rod cells. Proc Natl Acad Sci USA. 89(15):6841–6845.

Lamb TD, Hunt DM. 2017. Evolution of the vertebrate phototransduction cascade activation steps. Dev Biol. 431(1):77–92.

Lamb TD. 2013. Evolution of phototransduction, vertebrate photoreceptors and retina. Prog Retin Eye Res. 36:52–119.

Li W, Chen S, DeVries SH. 2010. A fast rod photoreceptor signaling pathway in the mammalian retina. Nat Neurosci. 13(4):414–416.

Liu Y, Zhou Q, Wang Y, Luo L, Yang J, Yang L, Liu M, Li Y, Qian T, Zheng Y, et al. 2015. *Gekko japonicus* genome reveals evolution of adhesive toe pads and tail regeneration. Nat Comm. 6(1):10033.

Marcellini D. 1977. Acoustic and visual displays behavior of Gekkonid lizards. Amer Zool. 17:251–260.

Menaker M, Moreira LF, Tosini G. 1997. Evolution of circadian organization in vertebrates. Brazil J Med Biol Res. 30:305–313.

Miller MA, Pfeiffer W, Schwartz T. 2010. Creating the CIPRES Science Gateway for inference of large phylogenetic trees in Proceedings of the Gateway Computing Environments Workshop. 14 Nov. 2010, New Orleans, LA. 1–8.

Okano T, Kojima D, Fukada Y, Shichida Y, Yoshizawa T. 1992. Primary structures of chicken cone visual pigments: vertebrate rhodopsins have evolved out of cone visual pigments. Proc Natl Acad Sci USA. 89:5932–5936.

Pedler C, Tilly R. 1964. The nature of the gecko visual cell. A light and electron microscopic study. Vision Res. 4:499–510.

Pinto BJ, Card DC, Castoe TA, Diaz Jr. RE, Nielsen SV, Trainor PA, Gamble T. 2019. The transcriptome of the Veiled Chameleon (*Chamaeleo calyptratus*): a resource for studying the evolution and development of vertebrates. Dev Dyn. 2019:1–7.

R Core Team. 2008. R: A language and environment for statistical computing. R Foundation for Statistical Computing. Vienna, Austria.

Revell LJ. 2012. phytools: An R package for phylogenetic comparative biology (and other things). Methods Ecol Evol. 3:217–223.

Röll B. 2000. Gecko vision: visual cells, evolution, and ecological constraints. J Neurocytol. 29:471–484.

Röll B. 2001. Gecko vision—retinal organization, foveae and implications for binocular vision. Vision Res. 41(16):2043–2056.

Roth LSV, Kelber A. 2004. Nocturnal colour vision in geckos. Proc R Soc Lond. 271:S485–S487.

Roth LS, Lundstrom L, Kelber A, Kroger RH, Unsbo P. 2009. The pupils and optical systems of gecko eyes. J Vis. 9(3):27.

Schott RK, Muller J, Yang CG, Bhattacharyya N, Chan N, Xu M, Morrow JM, Ghenu AH, Loew ER, Tropepe V, et al. 2016. Evolutionary transformation of rod photoreceptors in the all-cone retina of a diurnal garter snake. Proc Natl Acad Sci USA. 113:356–361.

Schott RK, Van Nynatten A, Card DC, Castoe TA, Chang BS. 2018. Shifts in selective pressures on snake phototransduction genes associated with photoreceptor transmutation and dim-light ancestry. Mol Biol Evol. 35(6):1376–1389.

Schwenk K. 1993. Are geckos olfactory specialists? J Zool. 229:289–302.

Serb JM, Oakley TH. 2005. Hierarchical phylogenetics as a quantitative analytical framework for evolutionary developmental biology. Bioessays. 27:1158–1166

Simões BF, Sampaio FL, Jared C, Antoniazzi MM, Loew ER, Bowmaker JK, Rodriguez A, Hart NS, Hunt DM, Partridge JC, et al. 2015. Visual system evolution and the nature of the ancestral snake. J Evol Biol. 28(7):1309–1320.

Simões BF, Sampaio FL, Loew ER, Sanders KL, Fisher RN, Hart NS, Hunt DM, Partridge JC, Gower DJ. 2016. Multiple rod–cone and cone–rod photoreceptor transmutations in snakes: evidence from visual opsin gene expression. Proc R Soc Lond. 283(1823): 20152624.

Springer MS, Emerling CA, Fugate N, Patel R, Starrett J, Morin PA, Hayashi C, Gatesy J. 2016. Inactivation of cone-specific phototransduction genes in rod monochromatic cetaceans. Front Ecol Evol. 4:61.

Stamatakis A. 2014. RAxML Version 8: A tool for phylogenetic analysis and post-analysis of large phylogenies. Bioinformatics. 30(9):1312–1313.

Taniguchi Y, Hisatomi O, Yoshida M, Tokunaga F. 1999. Evolution of visual pigments in geckos. FEBS Letters. 445(1):36–40.

Tansley K. 1964. The gecko retina. Vision Res. 4:33–37.

Underwood, G. 1951. Reptilian retinas. Nature. Lond. 167, 183.

Underwood, G. 1954. On the classification and evolution of geckos. Proc Zool Soc Lond. 123: 469.

Underwood G. 1957. On lizards of the family Pygopodidae. A contribution to the morphology and phylogeny of the Squamata. J Morphol. 100(2), 207–268.

Underwood G. 1970. The Eye. In: Gans C, editor. Biology of the Reptilia. New York: Academic Press. p. 1–97.

Walls GL. 1934. The Reptilian Retina: I. A new concept of visual-cell evolution. Am J Ophthalmol. 17(10):892–915.

Walls GL. 1942. The vertebrate eye and its adaptive radiation. Bloomfield Hills (MI): Cranbrook Institute of Science.

Yokoyama S, Blow NS. 2001. Molecular evolution of the cone visual pigments in the pure rod-retina of the nocturnal gecko, *Gekko gekko*. Gene. 276(1):117–125.

Xiong Z, Li F, Li Q, Zhou L, Gamble T, Zheng J, Kui L, Li C, Li S, Yang H, Zhang, G. (2016). Draft genome of the leopard gecko, *Eublepharis macularius*. GigaScience. 5(1):47.

Zhang X, Wensel TG, Yuan C. 2006. Tokay gecko photoreceptors achieve rod-like physiology with cone-like proteins. Photochem Photobiol. 82:145–1460.

Zhao, QY, Wang Y, Kong YM, Luo D, Li X, Hao P. 2011. Optimizing *de novo* transcriptome assembly from short-read RNA-Seq data: a comparative study. BMC Bioinformatics. 12(14):S2.

