## Supplementary material for "Transcriptomic data support a nocturnal bottleneck in the ancestor to gecko lizards": N/A. (from another version of this preprint: version 1, 2019-04-13)

*Supplemental Materials*

**Supplemental Figure 1:** GNAT phylogeny illustrating the presence of a functional copy of GNAT1 present in the genomes of *Gekko japonicus* and *Eublepharis macularius* that isn’t expressed in the eye of geckos.

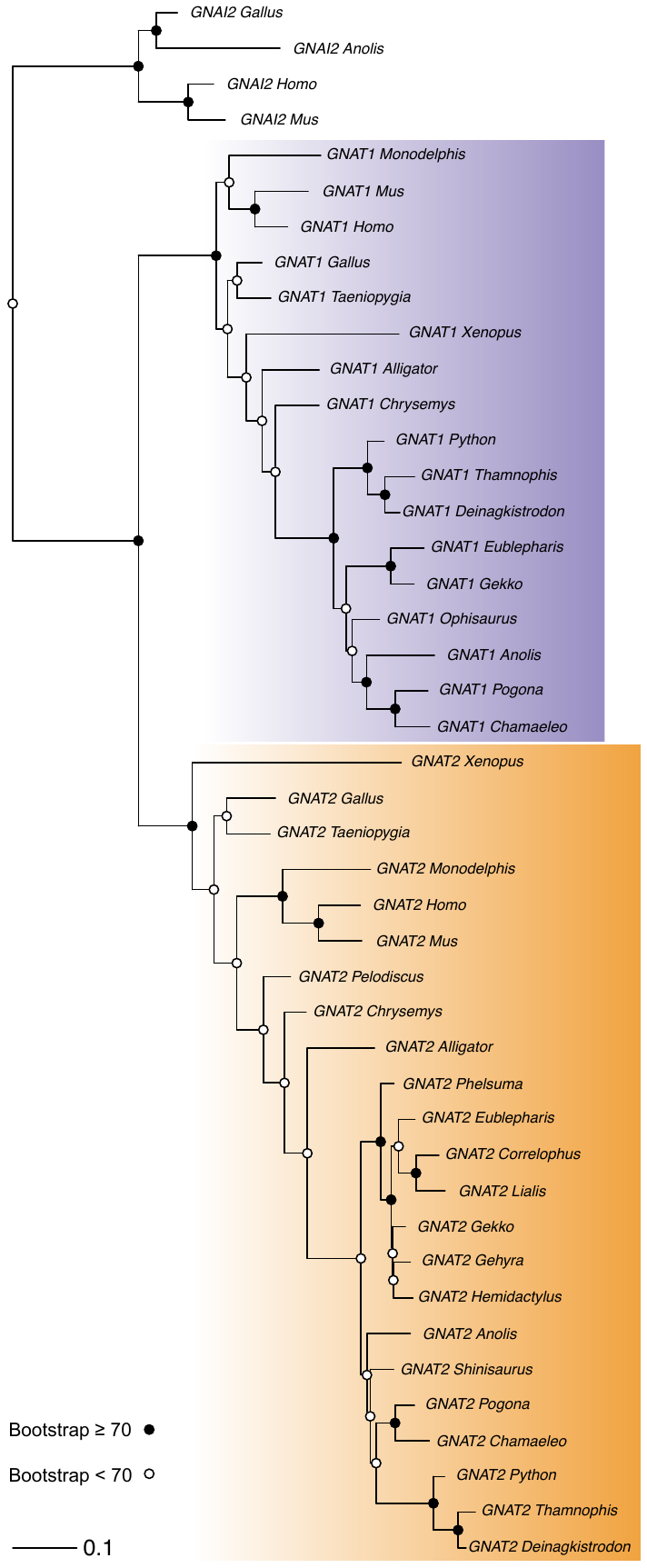

**Supplemental Figure 2:** PDE6 phylogeny illustrating the presence of PDE6C, not a PDE6 paralog, in geckos.

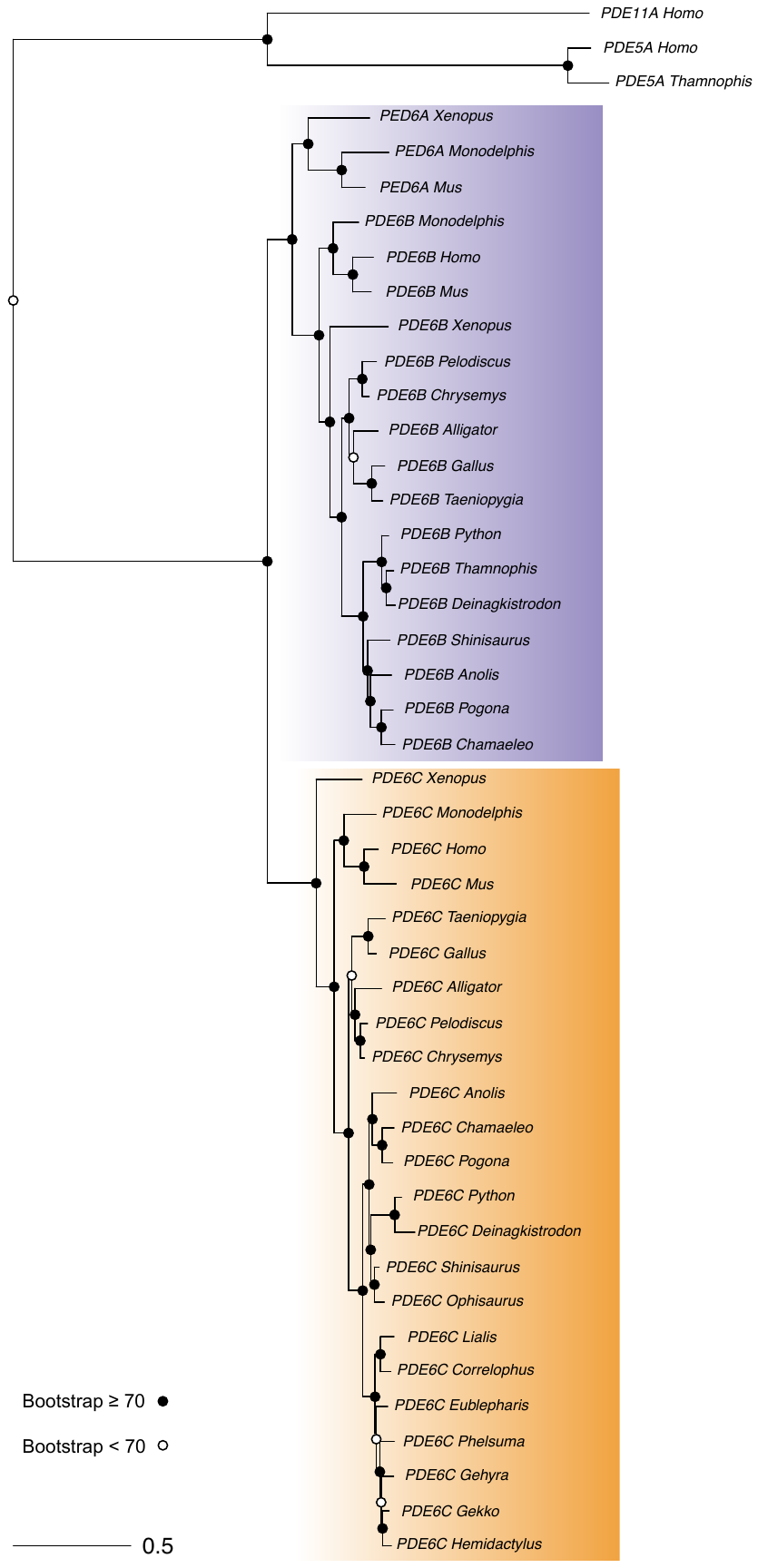

**Supplemental Figure 3:** CNG phylogeny illustrating the absence of CNGA1 in geckos.

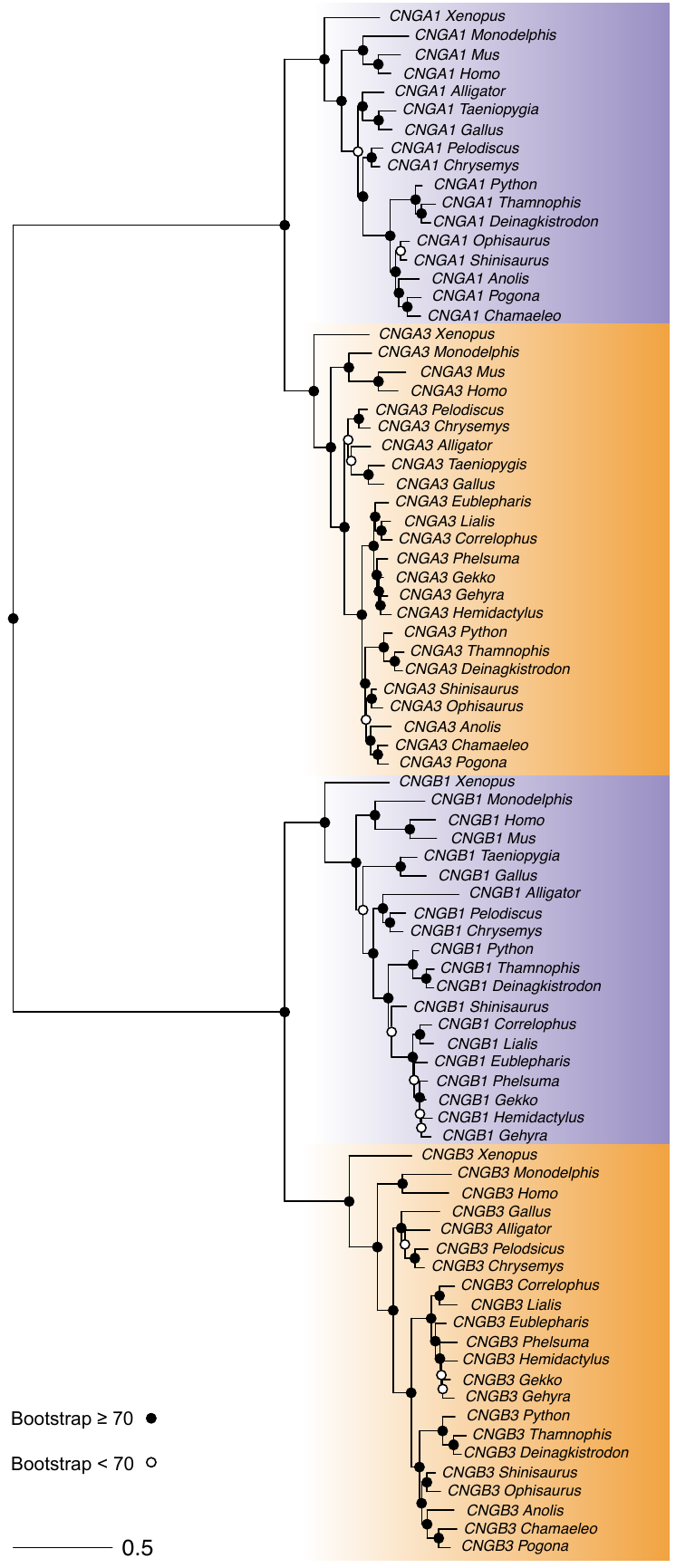

**Supplemental Table 1:** (part A) Sample information, read count, and number of transcripts assembled by DRAP. (part B) Assembly statistics, BUSCO and TransRate scores. (part C) number of transcripts successfully annotated and SRA accession information.

*part A:*

| **Species** | **Individual** | **Sex** | **Organ** | **Raw read pairs** | **Number of transcripts** |
| --- | --- | --- | --- | --- | --- |
| *Correlophus ciliatus* | TG2301 | M | eye | 17268534 | 130695 |
| *Gehyra mutilata* | TG2336 | M | eye | 27124576 | 73261 |
| *Hemidactylus turcicus* | TG2792 | F | eye | 34221650 | 71114 |
| *Lialis burtonis* | TG2108 | M | eye | 69110118 | 199317 |
| *Phelsuma laticauda* | TG3323 | F | eye | 68973597 | 232201 |
| *Chamaeleo calyptratus* | TG2597 + TG2872 | M + F | eye | 74120914 | 178294 |

*part B:*

| **BUSCO (%)** | **TransRate** | **N50** | **Number w/ ORF** | **GC Content** |
| --- | --- | --- | --- | --- |
| 82.6 | 0.1402 | 2142 | 40858 | 0.47 |
| 76.3 | 1.222 | 2279 | 26595 | 0.46 |
| 86.6 | 0.0952 | 2987 | 34566 | 0.46 |
| 88.6 | 0.158 | 1618 | 33621 | 0.45 |
| 89.4 | 0.1423 | 1295 | 34429 | 0.46 |
| 91.2 | 0.1155 | 2005 | 52470 | 0.44 |

*part C:*

| **# Annotated Transcripts** | **SRA Accession #** | **SRA Project #** |
| --- | --- | --- |
| 58591 | XXXXXX | XXXXXX |
| 38273 | XXXXXX | XXXXXX |
| 45643 | XXXXXX | XXXXXX |
| 64808 | XXXXXX | XXXXXX |
| 65910 | XXXXXX | XXXXXX |
| N/A. | XXXXXX | XXXXXX |

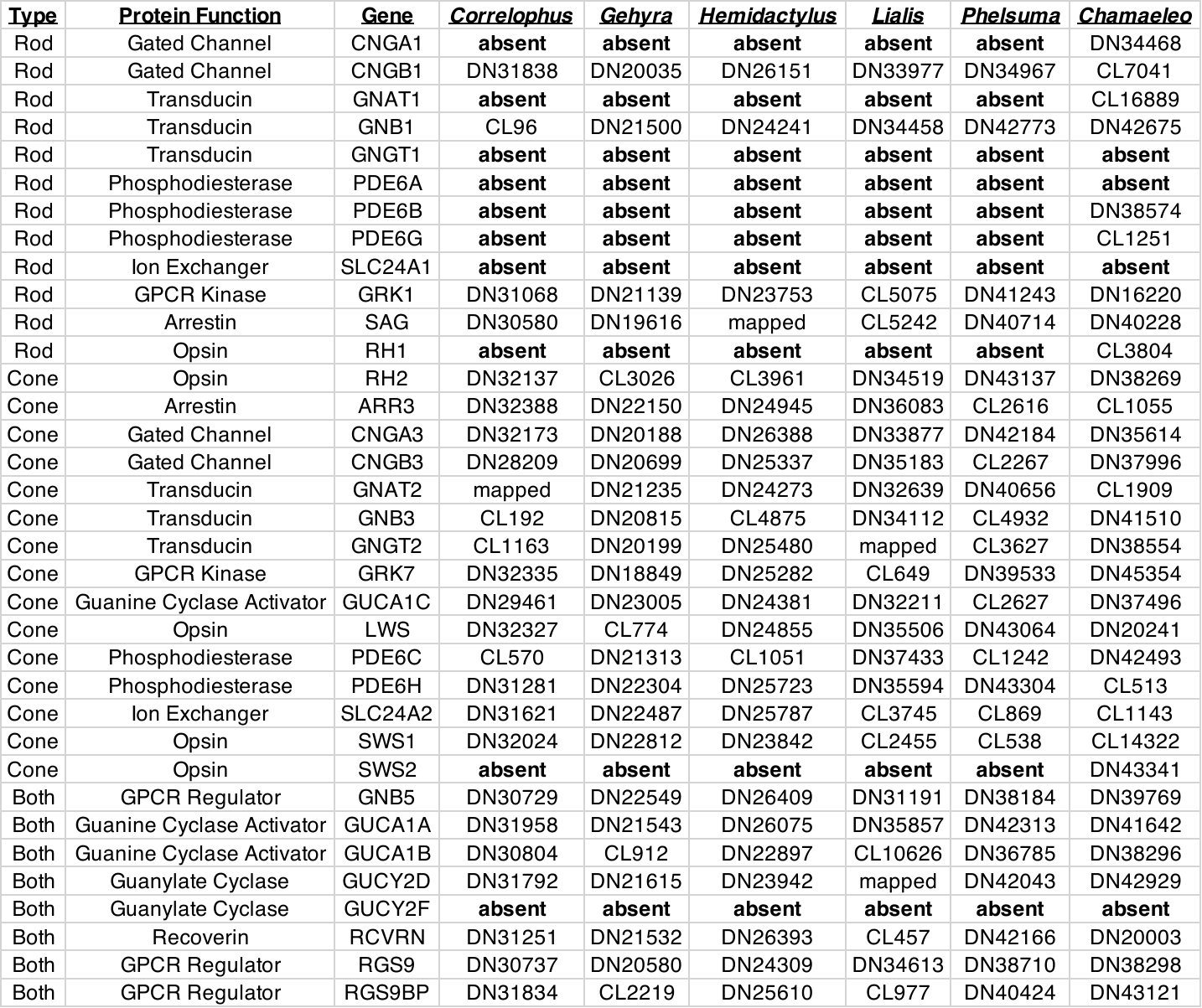
**Supplemental Table 2:** Presence/absence data for each of the 35 genes in the canonical phototransduction cascade. If present, the transcript ID is provided or is labeled as “mapped” clarifying that this transcript was not assembled *de novo*.
